# Aroma compounds profile is affected by the initial yeast ratio in wort co-fermentations

**DOI:** 10.1101/2024.03.01.583027

**Authors:** Jose Aguiar-Cervera, Federico Visinoni, Penghan Zhang, Katherine Hollywood, Urska Vrhovsek, Oliver Severn, Daniela Delneri

## Abstract

In recent years, the boom of the craft beer industry refocused the biotech interest from ethanol production to diversification of beer aroma profiles. This study analyses the fermentative phenotype of a collection of non-conventional yeasts and examines their role in creating new flavours, particularly through co-fermentation with industrial *Saccharomyces cerevisiae*. High-throughput solid and liquid media fitness screening compared the ability of eight *Saccharomyces* and four non-*Saccharomyces* yeast strains to grow in wort. We determined the volatile profile of these yeast strains and found that *Hanseniaspora vineae* displayed a particularly high production of the desirable aroma compounds ethyl acetate and 2-phenethyl acetate. Given that *H. vineae* on its own was a poor fermenter, we carried out mixed wort co-fermentations with a *S. cerevisiae* brewing strain at different ratios. The two yeast strains were able to co-exist throughout the experiment, regardless of their initial inoculum, and the increase in the production of the esters observed in the *H. vineae* monoculture was maintained, alongside with a high ethanol production. Moreover, different inoculum ratios yielded different aroma profiles: the 50/50 *S. cerevisiae*/*H. vineae* ratio produced a more balanced profile, while the 10/90 ratio generated stronger floral aromas. Our findings show the potential of using different yeasts and different inoculum combinations to tailor the final aroma, thus offering new possibilities for a broader range of beer flavours and styles.

**IMPORTANCE:** Craft brewing underwent an unprecedented growth in the last years due to customer demand for more unique and complex beverages. Brewers started to explore innovative fermentation methods using new ingredients, different brewing conditions, and new yeasts to explore a larger flavour landscape. The use of non-*Saccharomyces* yeasts has emerged as an effective strategy to produce novel distinct flavour profiles, however, knowledge regarding their fermentation performance and volatiles production is still limited, which hinders their industrial application. In this study, we expand on the knowledge of several non-*Saccharomyces* yeasts in terms of their brewing application and highlight the potential of *H. vineae* in co-fermentation with *S. cerevisiae* for producing unique fruity beers with a standard ethanol content. Our findings provide the craft beer industry with a new strategy to produce distinctive fruity beers.

## INTRODUCTION

The use of non-conventional yeasts (NCY) in the beverage industry has been rapidly growing in recent years (1). This is mainly due to the fact that these yeasts can produce beers with special characteristics, such as unique aroma profile (2–5) low ethanol content (6–8), probiotic properties (9, 10) and better nutritional content (11). Craft beer production has likewise flourished in the last decade: from 2012 to 2022 the number of active microbreweries in Europe has augmented a 73 % (12), and in contrast with large-scale industrial brewing, which relies on very well-established and standardized processes, craft brewing is more flexible in its approaches. The utilization of NCY in craft brewing represents a good opportunity for innovation to take place, yet the biological knowledge of NCY strains is still limited compared to the conventional brewing yeasts (*i.e. S. cerevisiae* and *S. pastorianus*).

High-throughput screening methods are a cost-effective approach to assess the suitability of conventional and non-conventional yeast strains for their use in brewing and have been previously employed for this purpose (5, 13). These methods enable the evaluation of numerous strains, aiding in the identification of those with desirable traits and reducing the number of candidates for downstream analytics of aroma compounds, ethanol, and other secondary metabolites such as glycerol and acetic acid. These compounds are key elements for any alcoholic beverage, as their concentration greatly influences the quality of the final product.

Higher alcohols such as isoamyl alcohol and phenylethyl alcohol, as well as their ester derivatives, isoamyl acetate and 2-phenylethyl acetate, are among the key aroma compounds that positively influence beer organoleptic quality, even at very low concentrations (14). These molecules are largely responsible for the fruity and floral notes that are highly esteemed in many beer styles, contributing to a complex and desirable flavour profile. On the other hand, certain volatiles including acetic acid, butyric acid, diacetyl, and phenolic compounds, are generally considered undesirable due to their tendency to produce off-flavours like sourness, rancidity, or buttery notes. However, concentrations of these compounds vary greatly within different beer styles; for instance, the smoky, clove-like character of phenolic compounds such as 4-vinyl guaiacol is essential in certain traditional ales, while low levels of diacetyl are desired in certain English ales and Bohemian Pilsners for the slight buttery note it imparts (15, 16). This complexity indicates the need for a nuanced approach to controlling these compounds in brewing, where the balance and concentration of each can either enhance or detract from the final product’s quality depending on the intended style and consumer preferences.

Besides optimizing fermentation parameters, namely growth rate, C/N ratio, oxygen, and temperature (17), aroma volatile compound production can be controlled through the use of genetically engineered yeast strains (17–21), however the use of genetically modified organisms (GMO) is forbidden in some parts of the world, such as Europe, where there is a strict GMO legislation. Moreover, some consumers can be sceptic of GMO products. An alternative and increasingly popular strategy involves the development of hybrid yeast strains. These hybrids are not classified as GMOs, and can produce higher levels of key aroma compounds (22–25), thus making their use in the food industry a feasible and attractive approach.

Co-fermentation between *Saccharomyces* and non-*Saccharomyces* yeasts represents yet another strategy for aroma enhancement. Typically, *Saccharomyces* sp. strains achieve robust fermentation performance, but produce low concentrations of certain desirable aroma compounds. Conversely, non-*Saccharomyces* yeasts, while less efficient in fermentation, can generate higher levels of key flavour compounds. By strategically combining conventional and non-conventional yeasts, brewers can harness the strengths of each, optimizing both fermentation efficiency and aroma. This approach has been extensively explored in the wine industry (26), and its use in brewing is also gaining traction in recent years.

Sequential and mixed beer co-fermentations of *S. cerevisiae* with non-*Saccharomyces* yeasts such as *Metschnikowia pulcherrima*, *Schizosaccharomyces pombe*, *Lachancea thermotolerans*, *Torulaspora delbrueckii* and several *Hanseniaspora* spp. and *Pichia* spp., have been trialled displaying very promising results (2, 4, 28–37). Here, we further expand on the potential application of non-*Saccharomyces* yeast in the brewing industry by analysing the growth, fermentation performance and aroma production of eight *Saccharomyces* and four non-*Saccharomyces* yeast strains in wort. Additionally, based on our analysis, we selected two promising strains (*S. cerevisiae* WLP095 and *H. vineae* Y-17530) for subsequent mixed wort co-fermentation at different ratios and assessed their population dynamics and aroma compound production (**Fig. 1**). We found that different inoculum ratios yielded different final flavour profiles, underlining the potential of tailoring the aroma of beer by means of different inoculation ratios in mixed co-fermentation.

**Figure 1.**
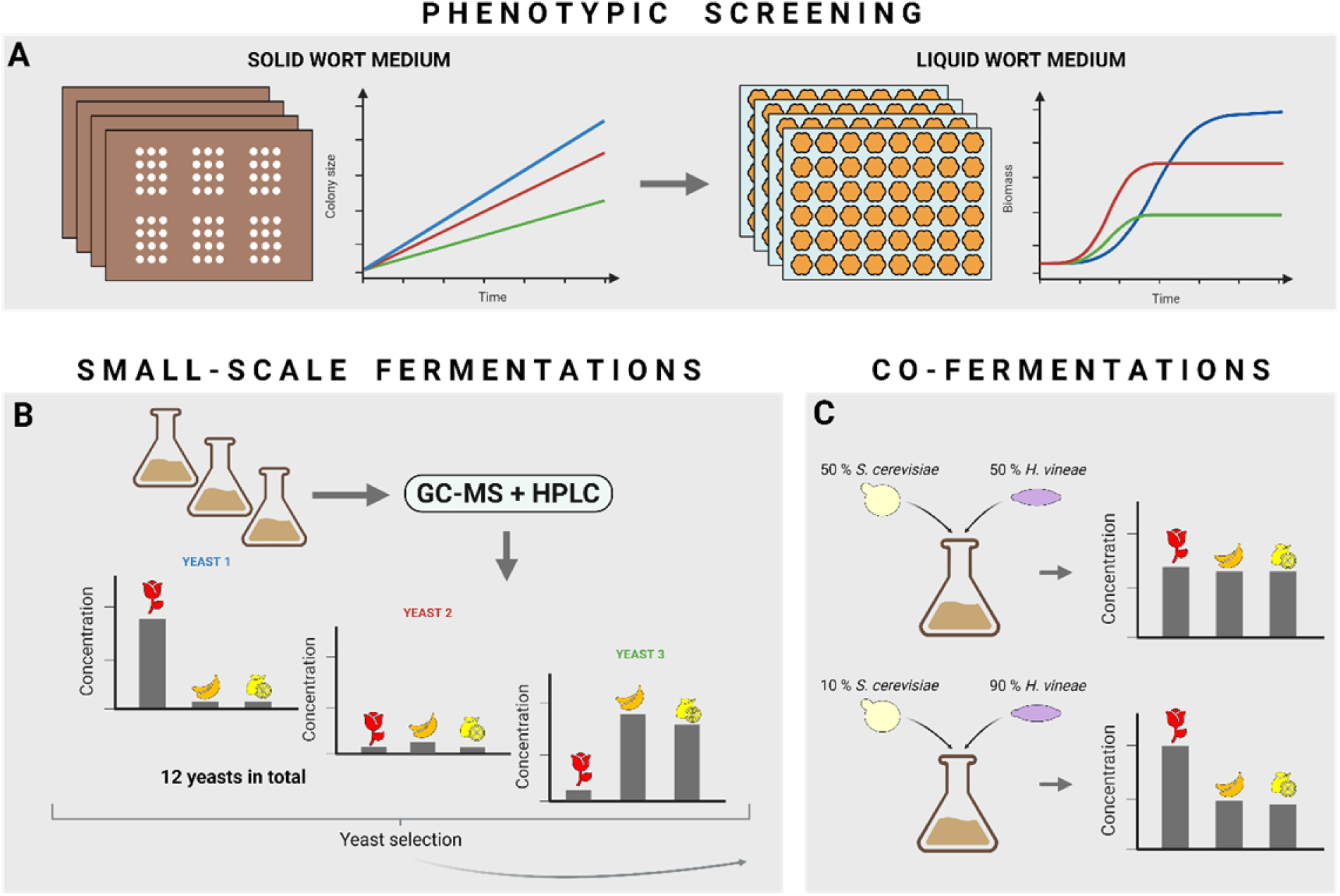
Experimental strategy: Firstly, we scored the ability of 12 yeast strains to grow in wort by screening their fitness in solid and liquid wort (**A**). Small-scale fermentations were then carried with 12 selected yeasts, and ethanol and aroma compound production (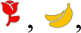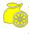) was assessed (**B**). Finally, based on our results, we selected two yeast strains for mixed wort co-fermentations using two different inoculum ratios and assessed the aroma profile (**C**).

## RESULTS AND DISCUSSION

### Phenotyping of yeast species grown on solid and liquid wort

With the aim of finding the ideal candidates for beer co-fermentation, we first screened the ability of 12 different *Saccharomyces* and non-*Saccharomyces* yeasts (**Table 1**) to grow in YPD and in beer-fermentative conditions consisting of 12 °Bx wort. It is key to ensure that any potential non-*Saccharomyces* candidate for beer co-fermentations is able to reach an acceptable biomass in wort to avoid out-competition when co-fermenting with a brewing *S. cerevisiae* strain.

**Table 1.**
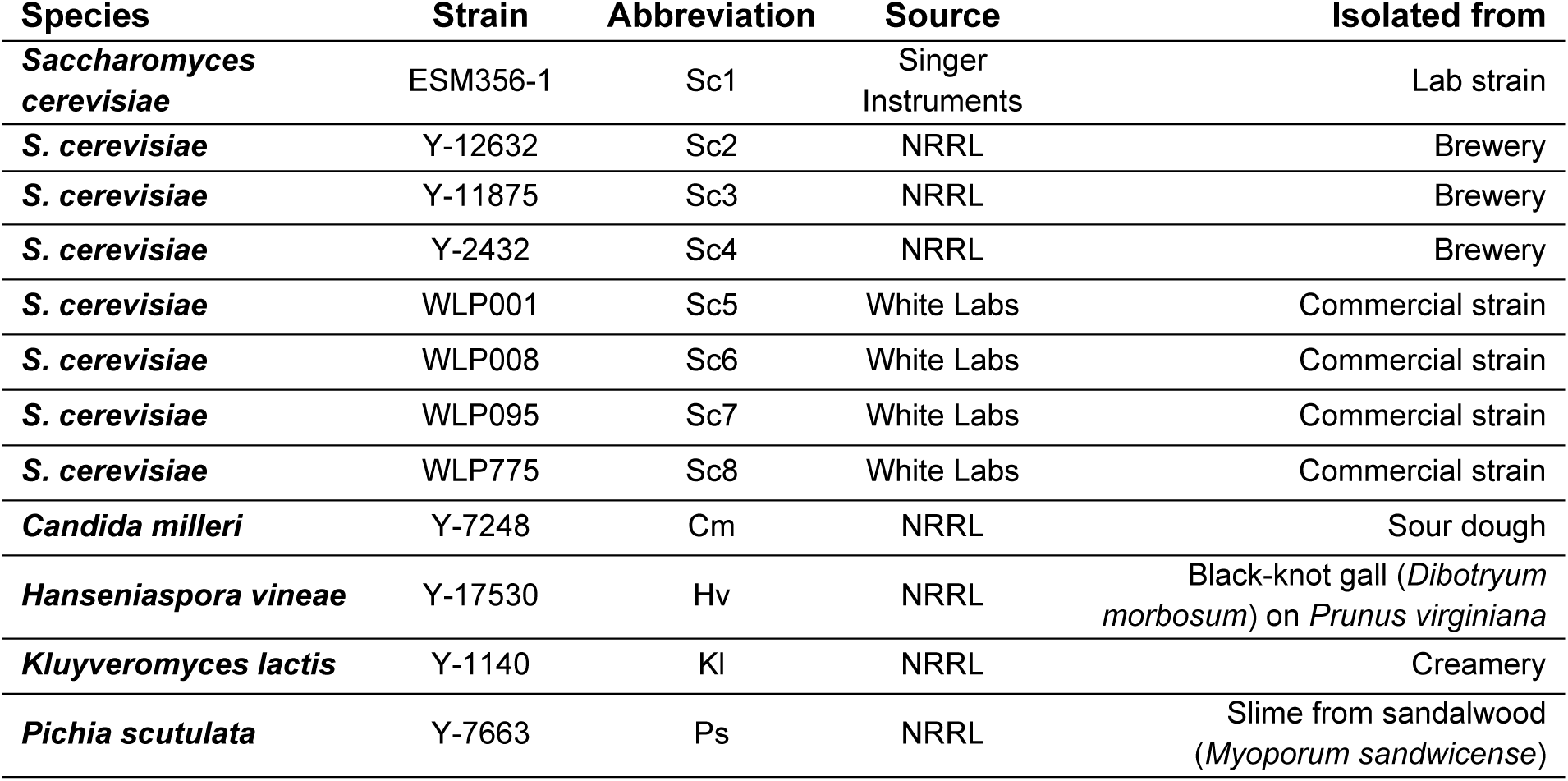
Yeast strains used in the study.

Phenotyping in solid media was carried out through adapting a high-throughput screening protocol that we have recently developed (13). This method uses colony size in solid media as a proxy for fitness.

We found that the fitness in wort for all the strains tested was either similar or worse to the fitness in YPD, with the exception of *S. cerevisiae* WLP008 (Sc6) which grew much better in wort. The NCY *Hanseniaspora vineae* Y-17530 (Hv) and *Kluyveromyces lactis* Y-1140 (Kl) displayed the biggest disparity of fitness, having final colony sizes in YPD that doubled the ones in wort. Overall, the results confirmed that all strains could grow using wort as a carbon source (**Fig. S1B**).

Next, micro-fermentations were carried out to assess the ability of the strains to grow in liquid wort, to mimic their potential performance in beer-fermentative conditions. This second screening also provided information regarding the fermentation performance in terms of sugar consumption and ethanol production. In this case, maximum growth rate (max. μ), maximum biomass, and Area Under the Curve (AUC) values were calculated for the 12 strains in both liquid YPD and wort after 48 h BioLector micro-fermentations at 20 °C.

In the experiments in liquid, the final biomass and AUC values were higher in YPD for all strains, including Sc6, however, the maximum growth rate was higher in wort for 9 out of 12 strains (**Tables 2 and 3**). This is rather intriguing since YPD has a higher glucose concentration than wort (20 g/L vs 12 g/L in wort) hence the expectation of higher growth rates in the medium with more glucose, which is the preferable carbon source for virtually any yeast. Here, it seems like the presence of other complex sugars such as maltose in wort, bolstered growth rate. On the other hand, YPD contains a lower C/N ratio than wort (37, 38), removing a growth limiting factor and allowing the accumulation of more biomass overtime.

**Table 2.** Growth kinetics parameters on the YPD (glucose 2 %) high-throughput micro-fermentations.

| Strain | Max. $\mu$ | Max. biomass | AUC |
| --- | --- | --- | --- |
| Sc1 | $0.24 \pm 0.02$ | $114.72 \pm 1.49$ | $3150.56 \pm 47.53$ |
| Sc2 | $0.32 \pm 0.04$ | $181.52 \pm 0.47$ | $6556.87 \pm 11.07$ |
| Sc3 | $0.29 \pm 0.01$ | $182.20 \pm 6.56$ | $6175.14 \pm 110.19$ |
| Sc4 | $0.29 \pm 0.00$ | $387.39 \pm 65.35$ | $9194.78 \pm 1094.25$ |
| Sc5 | $0.20 \pm 0.01$ | $182.20 \pm 6.56$ | $6175.14 \pm 110.19$ |
| Sc6 | $0.16 \pm 0.02$ | $207.37 \pm 0.53$ | $7490.57 \pm 12.65$ |
| Sc7 | $0.20 \pm 0.02$ | $185.15 \pm 0.48$ | $6688.01 \pm 11.30$ |
| Sc8 | $0.16 \pm 0.01$ | $185.85 \pm 6.69$ | $6298.64 \pm 112.39$ |
| Cm | $0.26 \pm 0.01$ | $336.11 \pm 37.68$ | $7984.90 \pm 428.30$ |
| Hv | $0.44 \pm 0.01$ | $677.60 \pm 5.31$ | $18702.96 \pm 440.35$ |
| KI | $0.27 \pm 0.02$ | $665.11 \pm 5.21$ | $19026.73 \pm 653.31$ |
| Ps | $0.29 \pm 0.03$ | $513.04 \pm 13.55$ | $12840.33 \pm 147.95$ |

**Table 3.**
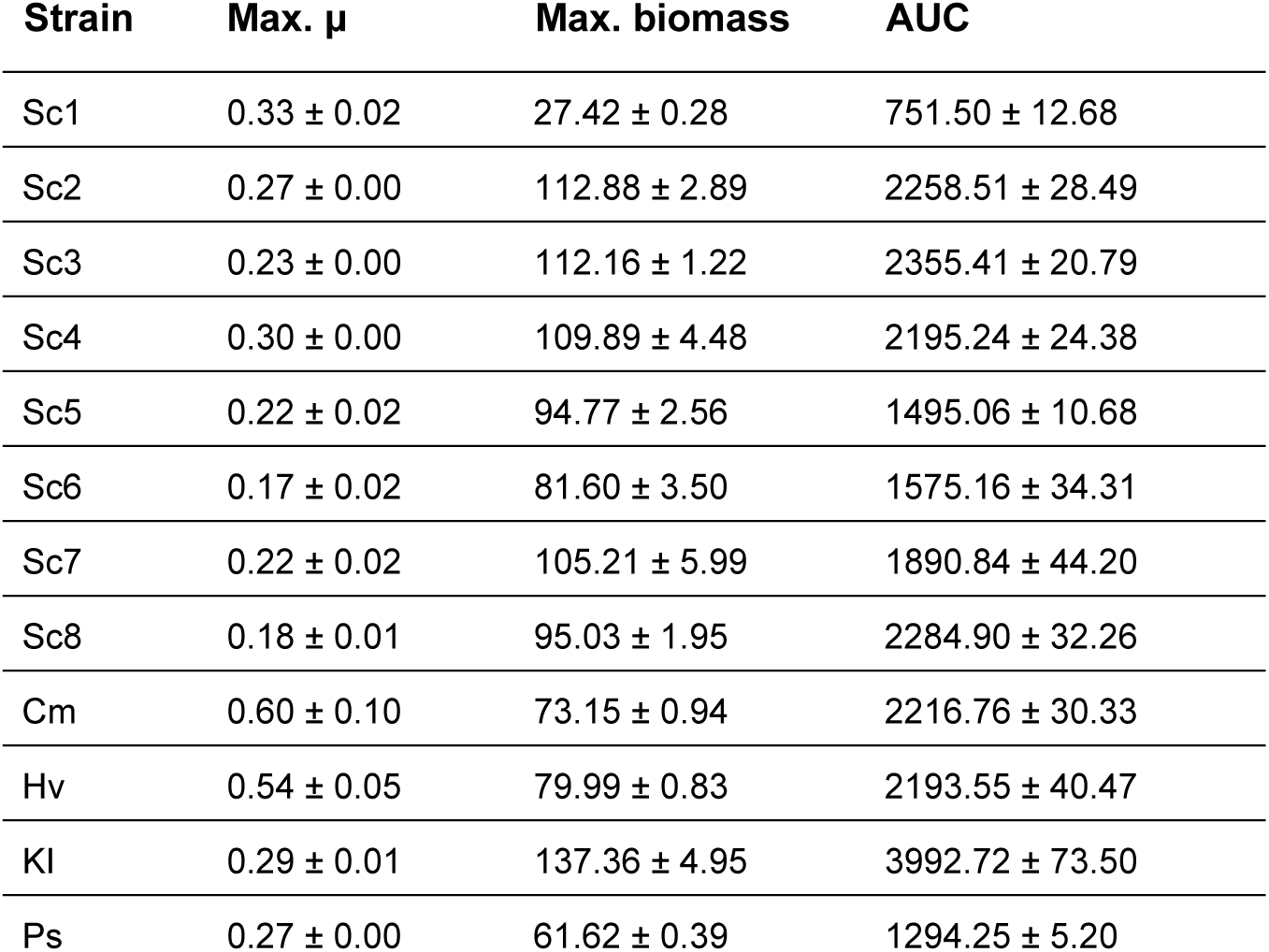
Growth kinetics parameters on the wort high-throughput micro-fermentations.

| Strain | Max. $\mu$ | Max. biomass | AUC |
| --- | --- | --- | --- |
| Sc1 | $0.33 \pm 0.02$ | $27.42 \pm 0.28$ | $751.50 \pm 12.68$ |
| Sc2 | $0.27 \pm 0.00$ | $112.88 \pm 2.89$ | $2258.51 \pm 28.49$ |
| Sc3 | $0.23 \pm 0.00$ | $112.16 \pm 1.22$ | $2355.41 \pm 20.79$ |
| Sc4 | $0.30 \pm 0.00$ | $109.89 \pm 4.48$ | $2195.24 \pm 24.38$ |
| Sc5 | $0.22 \pm 0.02$ | $94.77 \pm 2.56$ | $1495.06 \pm 10.68$ |
| Sc6 | $0.17 \pm 0.02$ | $81.60 \pm 3.50$ | $1575.16 \pm 34.31$ |
| Sc7 | $0.22 \pm 0.02$ | $105.21 \pm 5.99$ | $1890.84 \pm 44.20$ |
| Sc8 | $0.18 \pm 0.01$ | $95.03 \pm 1.95$ | $2284.90 \pm 32.26$ |
| Cm | $0.60 \pm 0.10$ | $73.15 \pm 0.94$ | $2216.76 \pm 30.33$ |
| Hv | $0.54 \pm 0.05$ | $79.99 \pm 0.83$ | $2193.55 \pm 40.47$ |
| KI | $0.29 \pm 0.01$ | $137.36 \pm 4.95$ | $3992.72 \pm 73.50$ |
| Ps | $0.27 \pm 0.00$ | $61.62 \pm 0.39$ | $1294.25 \pm 5.20$ |

In terms of the growth kinetics in YPD (**Table 2**), the *Saccharomyces* spp*. s*trains displayed max. μ values ranging from 0.16 to 0.32, whereas in the non-*Saccharomyces* spp., it ranged from 0.29 to 0.44, with Hv displaying the highest max. μ value. The major differences were observed in the max. biomass and AUC values, where the non-*Saccharomyces* spp. strains really stood up. For example, *C. milleri* Y-7245 (Cm) exhibited a max. biomass of 336.11 ± 37.68, which was the smallest of the non-*Saccharomyces* species but yet similar to the max. biomass of *S. cerevisiae* Y-2432 (Sc4) (387.39 ± 65.35), which was the best performer of the *Saccharomyces* genus. Hv and Kl displayed the highest. Max. biomass and AUC values, duplicating or even triplicating the values observed in the *Saccharomyces* strains. It is interesting to observe that the lab strain *S. cerevisiae* ESM356-1 (Sc1), which should be adapted to grow in lab conditions – namely YPD, did not thrive or surpass the non-lab strains in such condition. In fact, it displayed the lowest maximum biomass and AUC values of all the studied strains in YPD. This behaviour can be explained by the strain’s ploidy: Sc1 is haploid and the brewing *S. cerevisiae* strains tend to be polyploid (39). The presence of many copies of glucose transporter genes in the polyploid brewing strains will facilitate faster glucose intake and boost fitness.

Overall, the growth kinetics results in YPD confirmed that the non-*Saccharomyces* strains could generally grow as good as, or better, than the brewing *S. cerevisiae* strains.

When grown on wort, some of the non-*Saccharomyces* species displayed excellent maximum biomass and AUC values (**Table 3**). For example, Hv and Kl showed max. biomass values of 79.99 ± 0.83 and 137.36 ± 4.95, respectively. These values are similar and above the standard range we observed for the *S. cerevisiae* brewing strains (81.60 ± 3.50 – 112.88 ± 2.89). Cm showed a max. biomass value (73.15 ± 0.94) moderately lower than the *S. cerevisiae* brewing strains. Besides the lab strain (Sc1), *P. scutulata* Y-7663 (Ps) displayed the lowest max. biomass (61.62 ± 0.39) and AUC values in wort, so it was not deemed an ideal candidate for wort fermentation.

*C. milleri* is a species known for playing an important role in sourdough fermentations (40), an environment rich in sugars similar to the ones encountered in wort. Likewise, *H. vineae* is known to be widely present in wine fermentations (41), being well adapted to high concentrations of fermentation stressors such as ethanol. A good growth in beer-fermentative conditions for these two species was hence expected. *K. lactis* however, is not found in alcoholic fermentations but in dairy product fermentation. In fact, this yeast species is unable to grow in strictly anaerobic conditions and exclusively carries out respiration instead, mainly redirecting carbon into energy production and biomass anabolism at the expense of secondary metabolite production (42, 43). This physiological characteristic may explain the vigorous growth of Kl despite its lack of adaption for growing in wort.

In terms of fermentation performance, glucose was completely depleted in all cases, as expected, however, the consumption of maltose and maltotriose and the production of ethanol varied considerably between strains. Regarding YPD, the *S. cerevisiae* strains produced in between 5.50 ± 0.70 and 9.30 ± 0.14 g/L of ethanol, and the non-*Saccharomyces* species had a low fermentation performance, with the exception of Hv, which yielded 6.20 ± 1.13 g/L of ethanol (**Table S1**).

In wort, the strains that were able to metabolize the three main wort sugars (glucose, maltose and maltotriose), yielded, as expected, the highest ethanol concentrations. These were the brewing strains Sc4, *S. cerevisiae* WLP001 (Sc5), Sc6 and *S. cerevisiae* WLP095 (Sc7) **(Fig. 2A**, **Table S2**). The non-*Saccharomyces* yeasts had negligible ethanol production in wort due to their inability or deficiency to ferment maltose and maltotriose. Ps was able to metabolise slowly some of the maltose and maltotriose, producing 8.91 ± 0.76 g/L of ethanol. Partial utilization of maltose and maltotriose could be explained due to the presence of growth-limiting factors such as nitrogen depletion, ethanol stress or low pH. Another possible explanation would be that, as the micro-fermentations were not anaerobic (the plates were covered with a gas-permeable membrane, and aeration was active throughout the course of the micro-fermentations), at first the availability of oxygen would allow respiration to occur, boosting growth, but eventually, after biomass accumulation, the oxygen would become scarce and be the growth-limiting factor. High fermentation performance is considered an important requirement for traditional brewing, however, over the last decade, the demand for low-alcoholic beverages has been growing (8, 44); The use of low ethanol-producing non-conventional species has been previously proposed as an alternative for the production of low alcohol beer (6, 7, 45). The low-ethanol-yielding non-*Saccharomyces* yeasts of our study could be good candidates for the production of low alcoholic beer.

**Figure 2.**
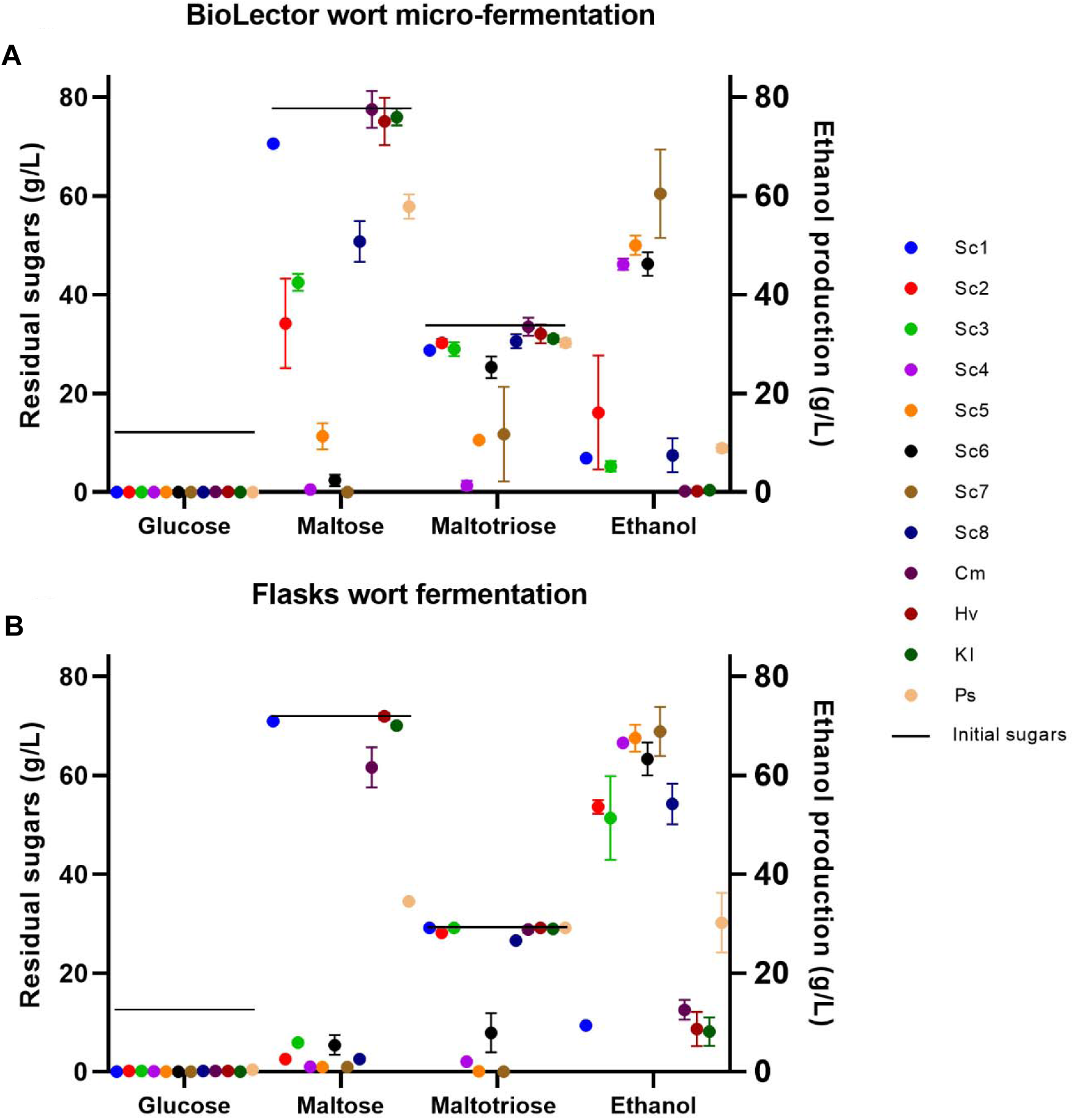
Residual sugars and ethanol production in the wort micro-fermentations (A) and small-scale flasks fermentations (B). Compounds were measured through HPLC at the end of the fermentations; initial wort sugars are represented with a straight line.

### Small-scale fermentations and flavour compound analysis

It is widely known that the size and the geometry of the fermentation vessel can have a profound impact on the fermentation performance mainly due to aeration and pressure (46, 47). So, to mimic real fermentation conditions more closely, we carried out wort small-scale fermentations in 500 mL flasks at 20 °C. Interestingly, ethanol production was significantly higher in the small-scale rather than in the micro-scale fermentations, especially in the case of *S. cerevisiae* Y-11875 (Sc2), *S. cerevisiae* Y-11875 (Sc3) and *S. cerevisiae* WLP775 (Sc8) (**Fig. 2B, Table S3**), likely due to the better consumption of maltose in the flasks experiment. This can be explained by the presence of oxygen in the micro-fermentations. Conversely, the small-scale fermentations were mainly anaerobic since the flasks were incubated statically and only de-caped for sampling. It is likely that the higher ethanol yield in the flasks was caused by a lower concentration of oxygen (48).

### Aroma compounds production

The production of nine key volatile compounds was measured through quantitative SPME GC-MS to help determine the ideal yeast combinations for co-fermentations (**Table S4**). Hv stood out as the strain with higher ethyl acetate (ethereal, fruity aroma) production (161.13 ± 72.22 mg/L), generating a concentration greatly surpassing its sensory detection threshold in beer (**Fig. 3, Table S4;** 50). The acceptable maximum concentration of ethyl acetate in wine is 200 mg/L (9), and the average value we observed for Hv is lower than this threshold. However, Hv’s production of ethyl acetate clearly surpasses the detection threshold seen in conventional lager beer (51), hence this would be a novel flavour trait, although we do not know yet how it would ultimately fare in different beer styles. This strain also produced the highest amounts of 2-phenylethyl acetate (floral, rose aroma; 2.77 ± 0.75 mg/L; **Fig. 3**, **Table S4**), and had a modest production of undesirable compounds such as acetaldehyde and 4-vinylguaiacol. This species has been previously reported to produce high levels of ethyl acetate and 2-phenylethyl acetate in the rich laboratory medium YPD and grape must (5, 52). Our results show that Hv maintains a similar production of those compounds also when grown in wort and would be a very good candidate for beer co-fermentations, where excessive production of those compounds should be limited.

**Figure 3.**
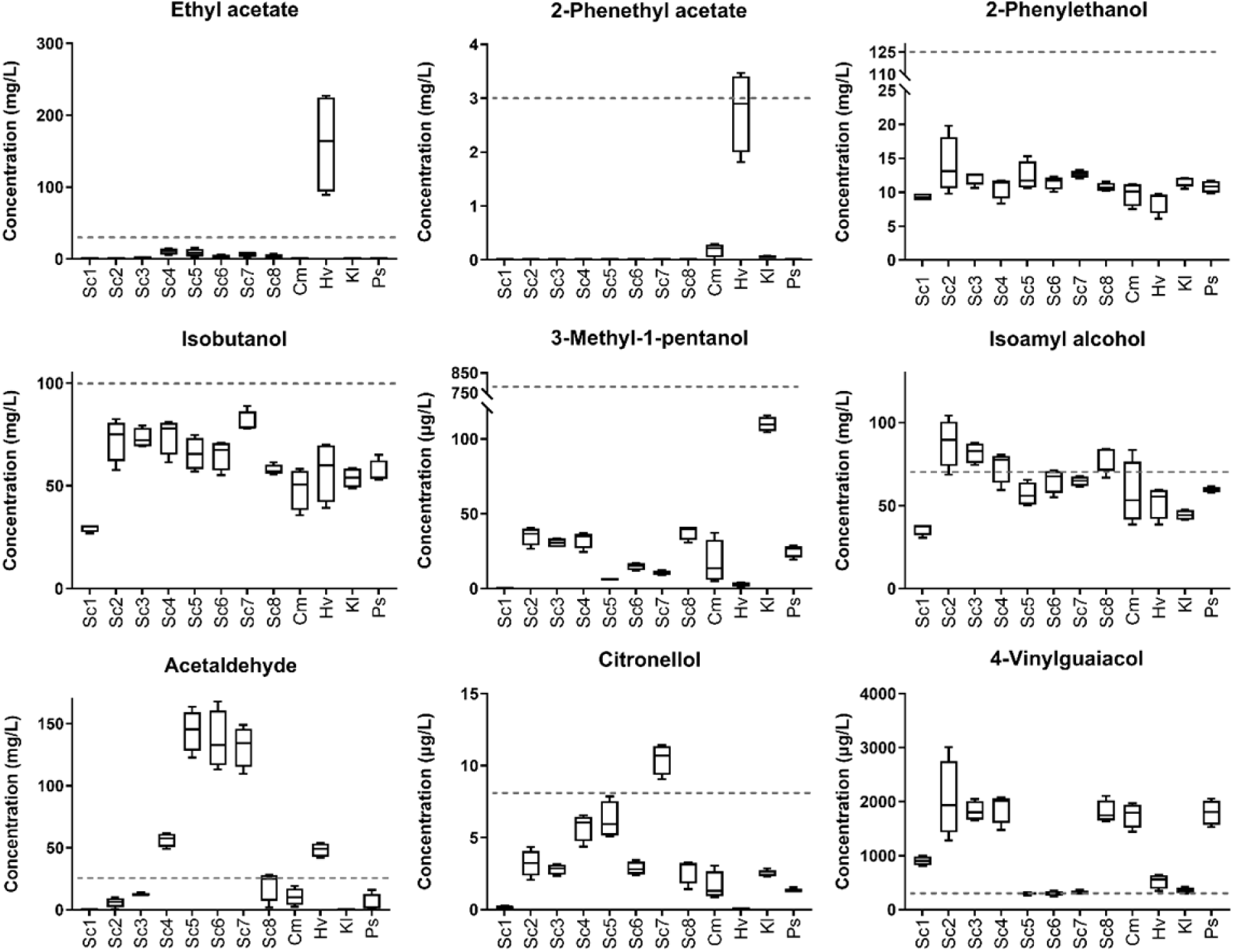
Concentration of key aroma compounds at the end of the wort flasks smallscale fermentation. Errors represent the standard deviation of the mean. The dashed lines represent the sensory detection threshold of each compound in beer.

The production of 2-phenyletylethanol (rose), isobutanol (sweet, solvent), and isoamyl alcohol (banana), was relatively similar in all the strains, and only in the case of isoamyl alcohol the production reached values beyond its sensory detection threshold (50) for some of the *S. cerevisiae* strains (**Fig. 3**). Acetaldehyde (apple, greasy) production was significantly higher in Sc5, Sc6 and Sc7. 3-Methyl-1-pentanol (apple, whisky) was produced in values below the sensory detection threshold in all cases, even in Kl which has been reported to be a good aroma producer (5).

The production of citronellol (rose, citrous), which is a key terpene that provides a floral and citrous aroma at very low concentrations (53), was highest in Sc7. Lastly, 4-vinylguaiacol (smoky) was produced significantly above its detection threshold in 7 out of 12 of the strains (**Fig. 3**, **Table S4**).

Overall, the aroma compound production of the commercial *Saccharomyces* strains (Sc5, Sc6, Sc7 and Sc8) was similar, and within the non-*Saccharomyces* yeasts, Hv stood up as a producer of the highest concentrations of the desirable esters ethyl acetate and 2-phenetyl acetate.

### Co-fermentations in wort

After measuring the fitness, the fermentation performance and the aroma compound production for our yeast strains in wort medium, we selected Sc7 and Hv for subsequent wort co-fermentations. Sc7 displayed good growth kinetics, good fermentation performance, and produced the highest amount of citronellol. Hv, despite its low fermentation performance, displayed high growth rate and biomass in wort and produced high amounts of ethyl acetate and 2-phenylethyl acetate. In the co-fermentations, the low fermentation performance of Hv would be compensated by the high fermentation performance of Sc7.

Prior to the co-fermentation experiments, different ratios of competitor strains were tested to find the optimum inoculum condition that allows maintenance of both strains in the cultures over time (**Fig. S2**). Interestingly, we observed that Hv coped well with the presence of Sc7 in all the strain ratios, and out-competition did not take place in any case. This is remarkable since more out-competition by Sc7 was expected, as is commonly observed in *Saccharomyces* non-*Saccharomyces* co-fermentation (54). The unexpected resilience of Hv underscores its potential for wort co-fermentation.

We then carried out flask co-fermentations with the inoculation ratios of 50/50 Sc7/Hv and 10/90 Sc7/Hv. Monoculture controls were also included (*i.e.* 100% Sc7 and 100% Hv).

As observed in the ratio optimization experiment (**Fig. S2**), Hv and Sc7 happily co-existed throughout the co-fermentation. Sc7 displayed final CFU/mL values close to 1 x 10^8^ when fermenting alone and in the 50/50 ratio. Hv also displayed a similar behaviour when alone or in the 50/50 ratio, with CFU/mL around 1 x 10^7^. In the 10/90 Sc7/Hv ratio, Hv started dominating the fermentation, but after 250 h Sc7 started to prevail, yet Hv sustained high growth (**Fig. 4**). Hv seemed to grow better in the presence of an equal amount of Sc7 cells compared to its monoculture. Some symbiotic effects may be happening in the 50/50 Sc7/Hv co-culture, which would boost the growth of Hv. Although such an effect is difficult to assess and can be attributed to different exometabolites excreted in the medium, one could hypothesize that the conversion of maltose and maltotriose to ethanol by Sc7 has a positive effect on the growth of Hv by making the carbon on those sugars accessible for Hv. In fact, Hv is not able to utilise those sugars as a carbon source but can metabolise ethanol. Additionally, it is worth considering that *Hansienospora* spp. can tolerate as much as 10 % v/v ethanol (55), so an increase in the ethanol levels as the one that we observe in our study should not be a major stressor for Hv.

**Figure 4.**
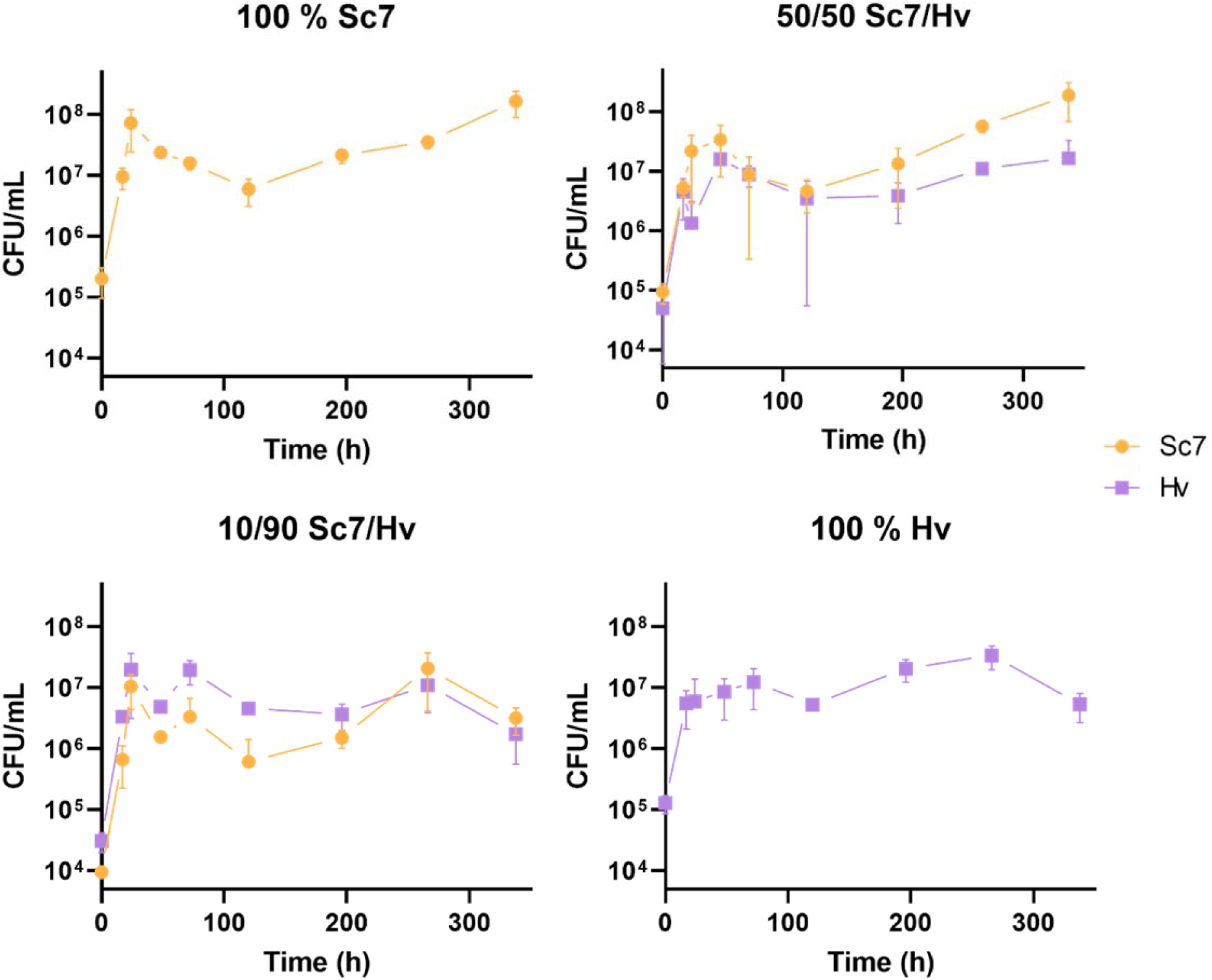
Population dynamics in the co-fermentations. Sc7 and Hv CFU/mL were tracked during the course of the wort co-fermentations to compare the growth of the co-fermenting yeasts and their monocultures.

In terms of fermentation performance, all sugars were completely depleted whenever Sc7 was present (**Fig 4**.). The consumption of sugars was virtually identical in both Sc7 monoculture and 50/50 Sc7/Hv co-fermentations (**Fig. 5**). In the 10/90 Sc7/Hv co-fermentation there was a delay in the depletion of maltose, and the final concentration of ethanol was lower by a very small margin (**Fig. 5; Table S5**). In any case, the production of ethanol in the 10/90 Sc7/Hv co-fermentation was more than sufficient for what is expected in a normal beer product, with a value of 50.23 ± 0.47 g/L. In the Hv monoculture, only the glucose was depleted, and the ethanol production was very low.

**Figure 5.**
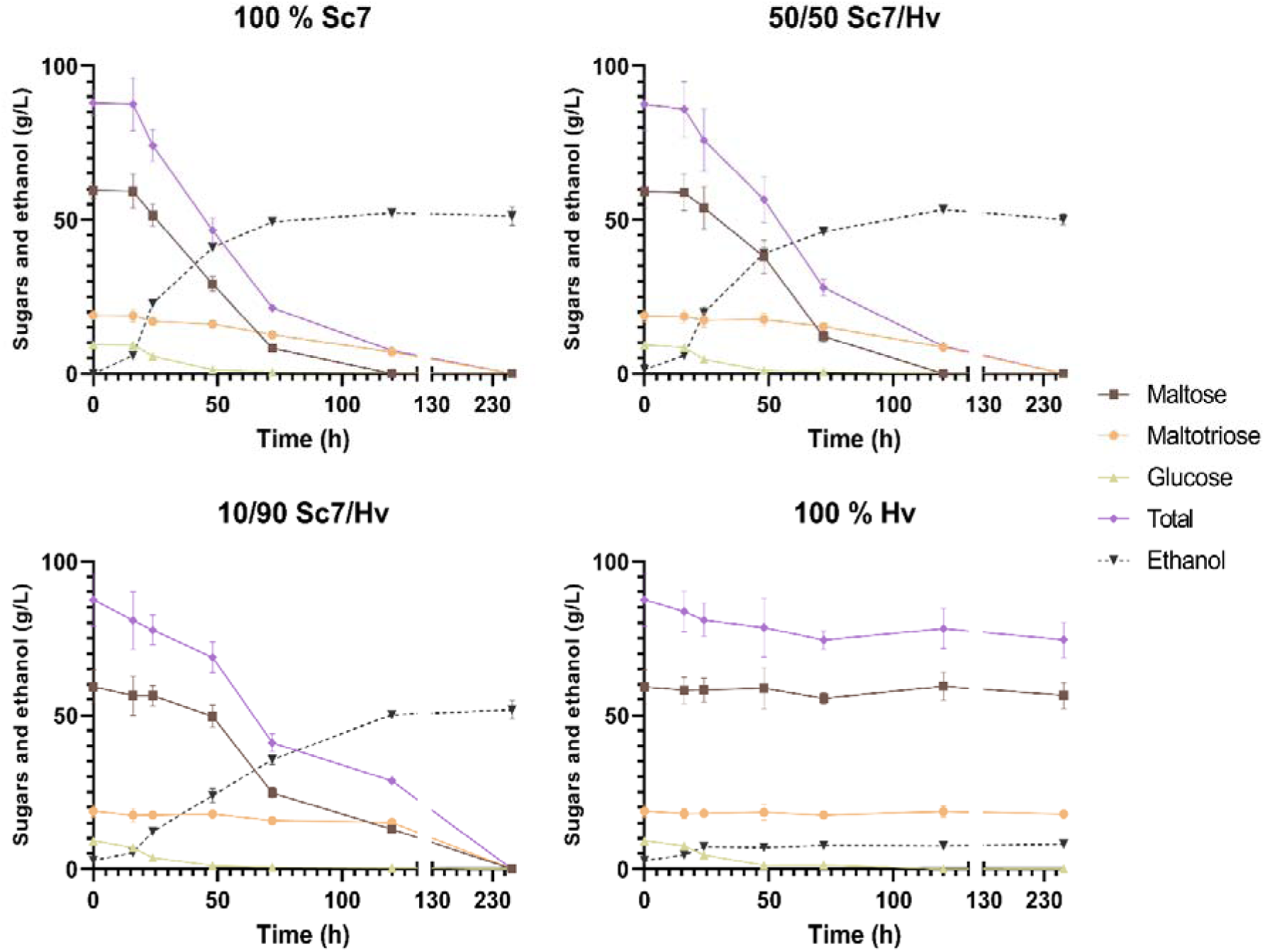
Sugar consumption and ethanol production during the Sc7 and Hv co-fermentations. Error bars represent the standard deviation of the mean.

Sequential beer co-fermentation, first with *H. vineae* and then with *S. cerevisiae* results in a large reduction in ethanol production compared to the *S. cerevisiae* monoculture (30), however, this was not observed in our study, which used mixed, simultaneous co-fermentations. This highlights the possibility of employing a mixed starter Sc7/Hv co-culture for brewing, instead of sequential co-fermentation, which is a more complex industrial procedure. The production of glycerol was significantly higher in the Sc7 monoculture and in the co-fermentations than in the Hv monoculture, while the acetic acid production was positively correlated with the amount of Hv in the inoculum (**Fig. 6, Table S5**). Glycerol is considered to be a desirable compound that provides body to the product and buffers off-flavours. Typically, in a beer product, its concentration ranges from 1 to 3 g/L (56). In our study, a good amount of glycerol was produced when *S. cerevisiae* was present in the fermentation, even in the ratio 10/90 Sc7/Hv co-fermentation. On the other hand, acetic acid can affect the aroma profile, and a concentration above its detection threshold (0.07 – 0.20 g/L) (50, 51) can negatively impact the quality of commercial ales and lager beers. However, acetic acid concentrations as high as 1.60 g/L are found in other beer styles such as gueuze (56). Although we do not know the exact detection threshold of acetic acid in our wort, in our study its production was potentially above or on the upper limit of the detection threshold in the Hv monoculture (0.36 ± 0.02 g/L) and in the 10/90 Sc7/Hv co-fermentation (0.20 ± 0.01 g/L) respectively (**Table S5**). It is worth highlighting that the odour detection threshold of a volatile compound can vary significantly between different matrices due to interactions within different aroma compounds (57). The optimal way to assess the potential impact of acetic acid in the co-fermentations would be through panel testing.

**Figure 6.**
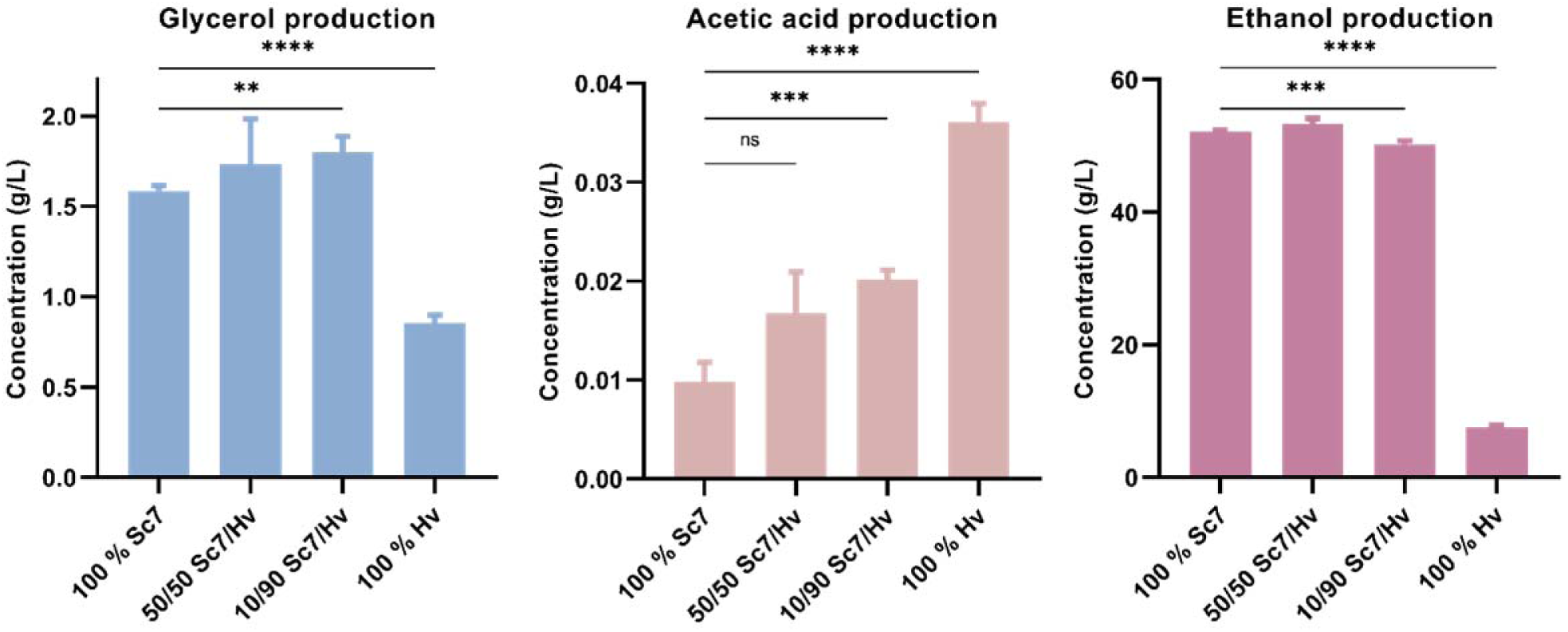
Glycerol, acetic acid and ethanol production in Sc7 and Hv wort co-fermentations. The graph represents the concentration of the compounds measured at the end of the fermentations. Error bars represent the standard deviation of the mean. Statistical differences were calculated through Tukey one-way ANOVA; **, adjusted p-value < 0.01; ***, adjusted p-value < 0.001; ****, adjusted p-value < 0.0001; ns, not significant.

### Aroma compound production in the co-fermentations

To assess the aroma profile of the co-fermentations, we carried out semi-quantitative SPME GC-MS on the end-product of the fermentations. We were able to identify twenty-one different flavour compounds relevant for brewing.

The presence of Hv increased the concentration of sixteen compounds associated with fruity and sweet descriptors, such as ethyl dodecanoate (waxy, sweet), 2-phenylethyl acetate (floral, rose), ethyl decanoate (waxy, sweet), ethyl 9-decenoate (fruity), ethyl acetate (fruity, ethereal), among others (**Fig. 7**; **Table 4**). 2-Phenylethyl acetate showed the highest increase when Hv was present, as observed in the small-scale monoculture experiments.

**Figure 7.**
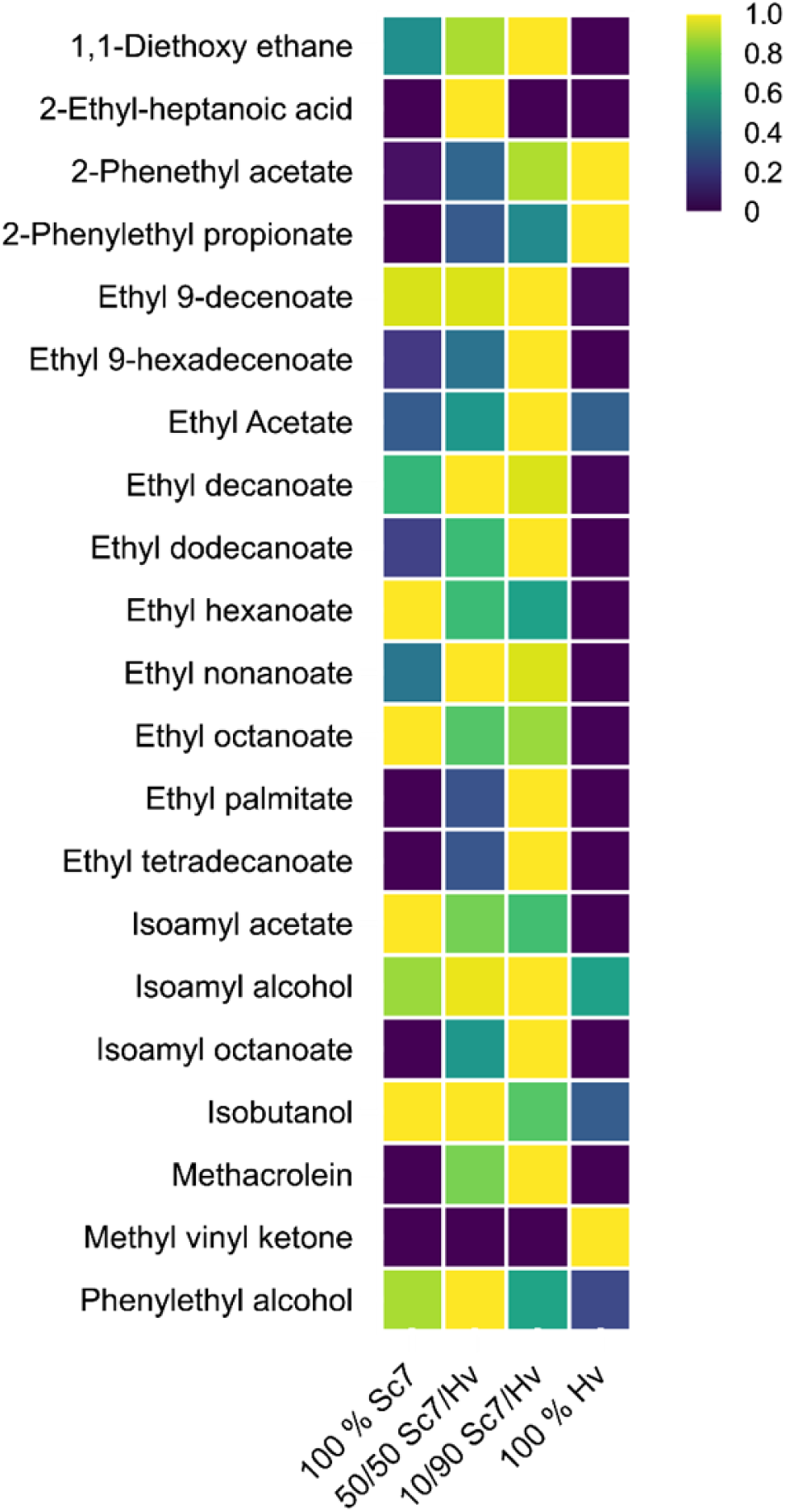
Heatmap of the aroma compounds measured in the co-fermentation. The plot represents the normalised relative area of the detected aroma compounds in the wort co-fermentations using different ratios of Sc7 and Hv.

**Table 4.**
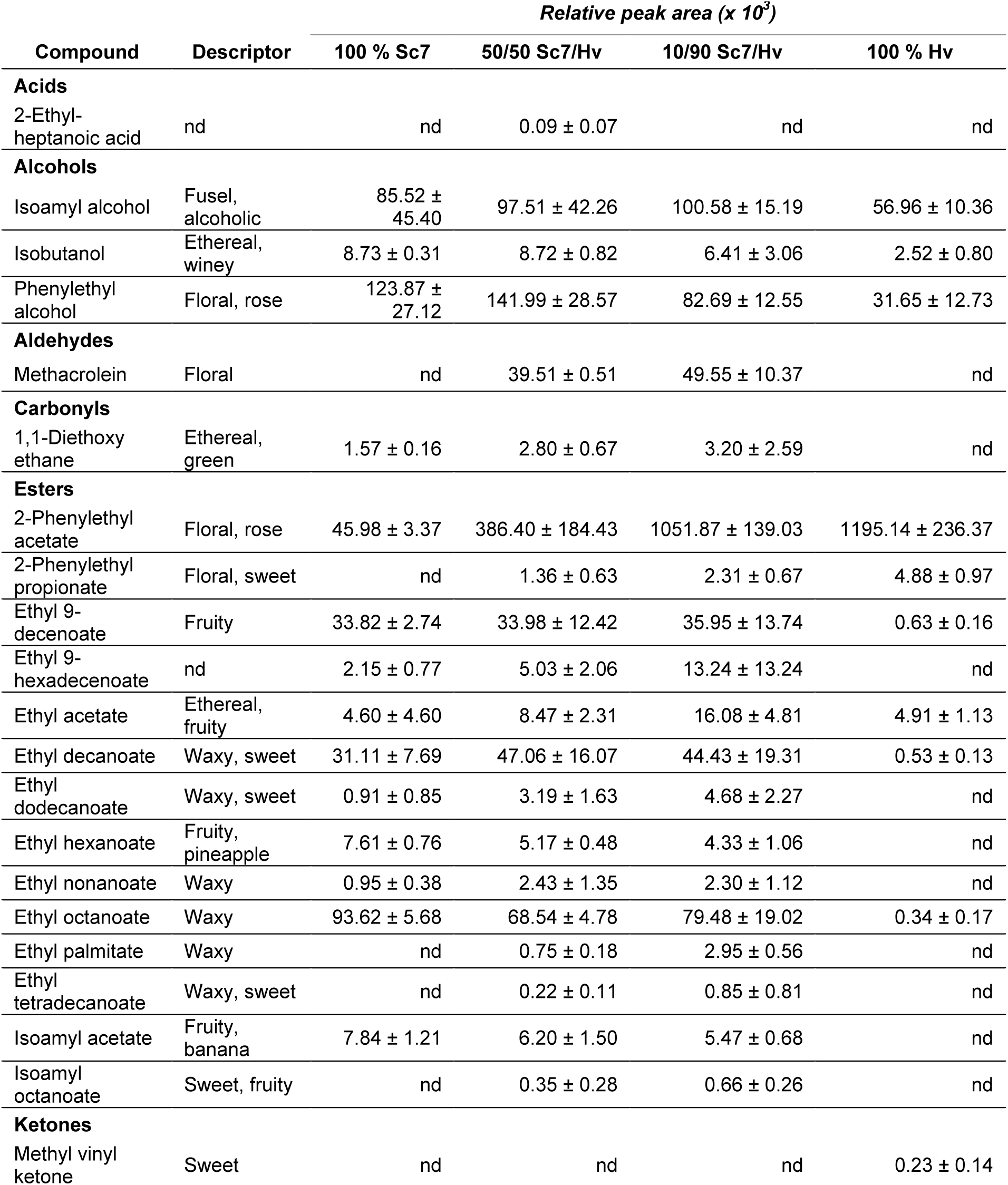
Relative area of the relevant aroma compounds detected in the co-fermentations. The relative areas were calculated through normalisation with an internal standard. Non-detected metabolites are represented with nd.

We classified the aroma compounds into three categories based on the production at the different ratios – *i*. compounds that showed the highest production in the Sc7 monoculture; *ii*. Compounds that showed the highest production in the Hv monoculture; *iii*. compounds that showed the highest production in the co-fermentations.

In the first category, we find isoamyl acetate (fruity, banana), a very important aroma in beer (16) with commercial *S. cerevisiae* brewing strains producing it in large quantities, likely due to domestication (39). Our wild Hv strain did not produce isoamyl acetate, but a relatively good amount was produced by Sc7, even in the 10/90 Sc7/Hv co-cultures. Ethyl hexanoate (fruity, pineapple) and ethyl octanoate (waxy, fruity) also fall in the first category, both compounds producing pleasant aromas (59, 60). Isobutanol (ethereal, winey) was also prevalently produced in the Sc7 monoculture, although Hv was also able to produce it to a small degree. This compound has been reported to decrease the aroma quality in wine (60) and could potentially impact the aroma profile of beer as well, although a very low concentration might be desirable. The second category includes 2-phenylethyl propionate, 2-phenylethyl acetate and methyl vinyl ketone. 2-Phenylethyl propionate (floral, sweet) is the result of the esterification of phenethyl alcohol and propionic acid, however, its specific contribution to alcoholic beverages quality has not been extensively explored in the literature. 2-Phenylethyl acetate, which is the result of the esterification of acetic acid and phenylethyl alcohol (61), provides a pleasant floral aroma and it is known to be highly produced in the *Hanseniaspora* sp. (4, 52, 63). We also detected this compound in the Sc7 monoculture, however, in the Hv monoculture and in the 10/90 Sc7/Hv co-culture its concentration was 2.5-fold and 2.3-fold higher, respectively (**Table 4**). The overproduction of this acetate ester has been recently attributed to the presence of several putative alcohol acetyltransferase genes in Hv, that would boost the conversion of phenylethyl alcohol to 2-phenylethyl acetate (63). Lastly, methyl vinyl ketone (sweet), was only found in the Hv monoculture in relatively low amounts.

Remarkably, the majority of highly abundant volatile compounds fall under the third category (co-fermentations), with the 10/90 Sc7/Hv co-fermentation displaying the highest productions overall (**Fig. 7**). Some of the most relevant compounds in this category were ethyl 9-hexadecenoate, ethyl dodecanoate, ethyl nonanoate, ethyl palmitate, isoamyl alcohol, isoamyl octanoate, phenylethyl alcohol and ethyl acetate, the latter displaying a concentration in the 10/90 Sc7/Hv co-culture which was three times higher than the one in the Sc7 monoculture (**Table 4**). Furthermore, it is interesting that compounds such as ethyl palmitate, isoamyl octanoate and methacrolein, were only found in the co-fermentations. This could occur due to a synergistic effect between both yeasts where the biosynthetic pathways to produce these metabolites lack one or more substrates, which are produced by the other co-culture yeast.

In general, a more balanced profile was observed in the 50/50 ratio, with a good production of virtually all the measured compounds without having any compound dominating. On the other hand, in the 10/90 ratio, we observed a higher production of 2-phenylethyl acetate and ethyl acetate. It is quite likely that a potential beer created with this Sc7/Hv yeast ratio would have a dominating floral and fruity character due to the high concentration of those two compounds, which can be desirable or not, based on consumer preference and beer style. Considering this, beer producers could tailor the aroma profile of the final beverage by using different inoculum ratios of Sc7 and Hv, creating a strong fruity beverage if using large ratios of Hv, or a more balanced one with a 50/50 ratio.

## CONCLUSIONS

We have confirmed that NCY have the potential to finely adjust the characteristics of beer by adding unique fruity and floral aromas. Our research has identified several NCY strains that display good growth performance as well as an enhanced aroma compound production in wort, Hv standing out as a high producer of ethyl acetate and 2-phenethyl acetate. The low ethanol production of these NCY opens two possibilities for their use in brewing: production of non-alcoholic beers, or co-fermenting with a high ethanol-yielding *Saccharomyces* sp.

We explored the latter approach and found that mixed co-fermentations employing 50/50 and 10/90 Sc7/Hv inoculation ratios significantly increased the overall production of desirable esters without compromising ethanol production compared to single-culture fermentations. Interestingly, certain esters like ethyl palmitate and isoamyl octanoate were only detected in the co-fermentation conditions suggesting a potential synergistic effect between these yeasts that requires further investigation. It’s worth noting that higher Hv inoculation ratios may result in increased acetic acid production indicating the importance of using lower Hv inoculation ratios to balance acetic acid levels in specific beer styles.

In conclusion, our study provides a basis for customising beer aroma profiles by manipulating the ratios of *S. cerevisiae* and *H. vineae* yeast ratios. This method provides brewers with the opportunity to tailor beers according to flavour preferences aligning with consumer demand for more complex and unique beverages. The results highlight the thrilling potential of yeast biotechnology in broadening the flavour range and sensory attractiveness of beers.

## MATERIALS AND METHODS

### Strains and media conditions

Twelve conventional and non-conventional yeast strains were employed in the study. The microbial strains used in this work were provided by the USDA-ARS Culture Collection (NRRL), Singer Instruments Co. Ltd., or obtained from White Labs. All the strains used in this study are listed in **Table 1**.

The media employed included 12 °Bx Spraymalt Amber 18EBC Wort (Brewferm Brouland, Belgium) and YPD (10 g/L yeast extract, 20 g/L peptone, 20 g/L glucose; Formedium, UK). The solidified media was supplemented with 20 g/L agar (Formedium, UK). A volume of 45 mL of media was poured into SBS PlusPlates (Singer Instruments, UK) and left to set on an even surface.

### High-throughput fitness assessment

An adapted version of our previously developed (13) colony-size-based fitness assessment was employed. The yeast strains were revived from –80 °C in YPD broth and arranged in 3×4 squares (**Fig. S1A**) on SBS PlusPlates containing solidified 12 °Bx wort and YPD using the PIXL (Singer Instruments, UK) robotic platform. The plates were replicated onto agar YPD and solidified wort plates using a ROTOR HDA (Singer Instruments, UK), sealed and incubated at 20 °C for 210 h. The PhenoBooth (Singer Instruments, UK) was used to capture images every 10 h. Following background subtraction, a colony radius value measured in pixels was obtained for each colony and time point and normalised using an in-house developed javascript script (https://github.com/joseac93/Normalisation-script.git), taking into account the plate, column and row means. The screening was performed in quadruplicate, including 12 in-plate replicates for each strain.

High-throughput micro-fermentations were carried out at 20 °C using a BioLector (m2p, Germany) on YPD and wort broth. Overnight YPD broth cultures of the 12 strains were used to inoculate the wells of a 48-well MTP-48-B FlowerPlate (m2p, Germany) containing fresh YPD or wort in a final volume of 1.5 mL and an OD_600_ of 0.1, in triplicate. The plate was sealed with a gas-permeable foil and incubated for 72 h with 800 rpm agitation, humidity control (>85% dH_2_O), and aeration (20.85 % O_2_). Scattered light values at values at 620 nm were measured automatically in each sample every 7.29 min.

### Small-scale fermentations

A scale-up experiment was performed in 500 mL flasks. Overnight YPD cultures of the 12 strains were used to inoculate 250 mL of wort at an OD_600_ of 0.1 in triplicate, then the flasks were covered with sterile cotton pads and incubated statically at 20 °C for five days, (until all metabolizable sugars were depleted). Samples for downstream analysis were collected from each flask every 24 hours.

### Co-fermentations

To identify the optimal inoculation ratio of Sc7 and Hv to avoid out-competition, five wort co-fermentations and two monocultures, one for each species, were set up in triplicate in a 96-deep-well plate. The medium was inoculated to a final combined OD_600_ of 0.1 employing the following ratios: 100 % Sc7, 50/50 Sc7/Hv, 30/70 Sc7/Hv, 20/80 Sc7/Hv, 10/90 Sc7/Hv, 5/95 Sc7/Hv and 100 % Hv. The deep-well plate was covered with an aluminium seal and incubated statically at 20 °C.

The growth dynamics were studied by comparing the CFU/mL for each yeast species on each ratio. Colonies of Sc7 and Hv were easily distinguishable by eye due to their colony morphology characteristics (**Fig. S3**): Sc7 formed very bubbly and thick colonies, whereas Hv formed thinner colonies, which were surrounded by a halo of less dense colony material and were slightly raised at the centre. Samples for CFU assessment were taken at 0, 12, 24, 48, 72, and 96 h.

Co-fermentations were performed in 500 mL flasks, in triplicate. Cell count was carried out with a Cellometer Auto M10 (Nexcelom Bioscience, USA) to establish the initial inoculum. Two hundred and fifty mL of 12 °Bx wort were inoculated with a final concentration of cells of 1 x 10^5^ cells/mL in the following ratios: 100 % Sc7, 50/50 Sc7/Hv, 10/90 Sc7/Hv, and 100 % Hv.

The flasks were covered with sterile cotton pads and incubated statically at 20 °C. Samples were taken at 0, 17, 24, 48, 72, 120, 192, 262 and 334 h. Population dynamics was studied with the same CFU method as the one employed in the pre-study, based on colony morphology differences.

### HPLC and GC/MS analyses

A 1260 Infinity II LC High-Performance Liquid Chromatography system equipped with a Refractive Index Detector (Agilent, USA) and a 300×7.8 mm Hi-Plex Exchange column (Agilent, USA) was used to quantify glucose, maltose, maltotriose, ethanol, glycerol and acetic acid at the end of the BioLector micro-fermentations and at different timepoints during the course of the flasks small-scale fermentations and co-fermentations. H_2_SO_4_ 5 mM was utilized as solvent with a flow rate of 0.8 mL/min at 55 °C. The compounds were detected by retention times and quantified using calibration curves made with analytical grade standards (Sigma-Aldrich, Germany).

A Thermo Scientific TSQ Quantum GC Triple Quadropole GC-MS was used to quantify the volatile flavour compounds in the small-scale fermentations. Twenty-five µL of internal standard (2-octanol 2.5 mg/L) and 0.5 g of NaCl were added to each sample in 20 mL vials. After 10 min of incubation at 40 °C, volatile compounds were collected on a Divinylbenzene/Carboxen/Polydimethylsiloxane 2 cm fibre (DVB-CAR-PDMS Supelco, USA) with an extraction time of 30 min. A VF-wax column (Agilent, USA) 30 m/I.D 0.25 mm/Film 0.25 μm was used for the separation. Oven temperature was set at 40 °C for 4 min and then increased 6 °C / min until 250 °C were reached. The final temperature was kept for 5 min. The injector and interface temperatures were kept at 250 °C as well. Helium was used as the carrier gas with a flow rate of 1.2 mL/min. The time for thermal desorption of analytes was 4 min. The MS detector was operated in full scan mode at 70 eV with a scan range from 40 to 300 m/z for 44 min.

Quantitative data analysis was performed using ThermoXcalibur (Version 2.2 SP1.48, Thermo scientific, USA). Identification of compounds was based on comparison with a mass spectral database (NIST version 2.0) and with pure reference standards. The calibration curves were done by adding the standards in a MQ water + 5 % EtOH solution (**Fig. S4**).

The relative concentrations of the volatile compounds of the co-fermentations samples were assessed using a Gerstel MPS dual head system (Gerstel, Germany) and an Agilent 7890B Gas Chromatograph (GC) in conjunction with an Agilent 5977B Series Mass Selective Detector (MSD) (Agilent, USA). Five mL of each sample were placed in a 20 mL vial with 0.5 g of NaCl and 25 µL of the internal standard (2-octanol 2.5 mg/L). The samples were then incubated for 10 minutes at 30 °C to reconcentrate volatile analytes into the headspace, after which a PDMS fibre (100 μm thickness) was extracted for five minutes and injected into the front inlet. An Agilent VF-5MS column (30 m x 25 mm x 0.25 μM) with a flow rate of 1 mL min-¹ was used to carry out the GC separation. The gradient in the oven was: hold at 40 °C for 4 minutes, a linear 6 °C per minute gradient to 250 °C, and a final hold at 250 °C for 5 minutes. The GC-MS analysis time for each sample was 44 min. The temperature conditions were as follows: inlet 280 °C, transfer line 300 °C, EI source 230 °C, quadrupole 150 °C. The MS scanned mass range was 40-400 m/z.

## ACKNOWLEDGEMENTS

This work was supported by the European Commission (H2020-MSCA-ITN-2017) grant number 764364 and by Biotechnology and Biological Sciences Research Council (BBSRC) grant number BB/R000069/1 awarded to DD. We also would like to acknowledge the USDA-ARS Culture Collection (NRRL) for providing microbial strains and the Faculty of Science and Engineering Mass Spectrometry and Separation Facility (RRID: SCR_024761) for support.

